# Physiological state matching in a pair bonded poison frog

**DOI:** 10.1101/2022.09.25.509360

**Authors:** Jessica P. Nowicki, Camilo Rodríguez, Julia C. Lee, Billie C. Goolsby, Chen Yang, Thomas A. Cleland, Lauren A. O’Connell

## Abstract

More than a century ago, Charles Darwin hypothesized that the empathy-like phenotype is a phylogenetically widespread phenomenon. This idea remains contentious, due to the challenges of empirically examining emotions, and few investigations among non-mammalian vertebrates. We provide support for Darwin’s hypothesis by discovering partial evidence for the most ancestral form of empathy, emotional contagion (i.e., matching another individual’s emotional state), in the pair bonding mimetic poison frog, *Ranitomeya imitator*. We found that male corticosterone, a physiological biomarker of stress, positively correlates with female partners in experimental and semi-natural conditions. This does not appear to coincide with behavioral state-matching. However, it is specific to female partners relative to familiar female non-partners, and is independent of effects that commonly confound studies on emotional contagion. Furthermore, this physiological state-matching is irrespective of partnership longevity or lifetime reproductive output. These results physiologically indicate socially selective emotional contagion in a monogamous amphibian, and paradigms that elicit coinciding neural and behavioral indicators and morphogenic co-variation are needed for further corroboration. Further studies on ancestral forms of empathy in non-mammalian vertebrates are warranted.

## Introduction

Emotional contagion, described as the ability to match the emotional state of another individual, is considered the most fundamental and ancestral form of empathy (1).This contagion or “resonance” of emotion, relies on a simple perception-action mechanism where the perception of a “demonstrator’s” emotional state triggers a neurophysiological representation of the same emotional state (i.e., “state matching”) in the “observer” (2,3). Consequently, it often prompts adaptive behavioral responses such as risk avoidance, social cohesion, conflict resolution, and social bonding (1,3,4). More than a century ago, Charles Darwin provocatively argued that the empathy-like phenotype is not unique to humans, but rather is phylogenetically widespread, after observing behavioral signs of sympathetic distress across a variety of species (3,5). To date, the evolutionary origins and phylogenic scope of the empathy-like phenotype remain contentious (3,6), partially owing to the scarce empirical evidence beyond mammals and birds.

Emotional contagion may be especially apparent within pair bonds, a socio-sexual system emerging in humans and only 1-9% of other non-avian vertebrates (7). Pair bonded relationships display among the highest levels of key empathetic characteristics, including ingroup familiarity and selectivity, social closeness, and cooperation (3,8). For example, pair bonded individuals rely almost exclusively on each other to execute highly coordinated activities that are critical to inclusive fitness, including bi-parental care, joint territory defense, and reciprocal predator vigilance (9–12). Since the fitness of pair bonded individuals is heavily reliant on each other, a mechanism for appropriately understanding and attending to each other’s needs is critical. However, outside of humans, emotional contagion within pair bonded relationships has only been studied in voles and zebra finches (13,14), leaving open the possibility of this phenomenon in other pair bonding vertebrates.

Here, we tested for evidence of emotional contagion in a pair bonding amphibian, the mimetic poison frog (*Ranitomeya imitator*). In this species, males and females form prolonged partnerships characterized by affiliative interactions, the mutual defense of a shared territory, and joint care for offspring (15,16). By subjecting male-female partner dyads to an “empathy assay” similar to one developed for rodents (13), we tested the hypothesis that male observers would display emotional contagion and ingroup bias towards female partners (demonstrators) that were subjected to a stressor. Specifically, we predicted that males would state match female partners hormonally and behaviorally despite never experiencing or observing the stressor themselves, and that this response would be biased towards partners relative to familiar non-partner females. Furthermore, we predicted that this response would increase with the longevity and lifetime reproductive output of partnerships. To better interpret the results from the stress experiment and establish the semi-natural (baseline) corticosterone state of pairs, we also examined corticosterone levels and hormone matching of experimentally naïve pairs during cohabitation.

## Methods

### Establishment, longevity, and lifetime reproductive output of pair bonds

All procedures were approved by the Stanford University Animal Care and Use Committee (APLAC protocol numbers 32961 and 33880). *Ranitomeya imitator* used in this study were sexually mature adults reared in our breeding colony. Sub-adults were co-housed in group terraria until they reached the size of sexual maturity. Glass housing terraria (30.48 × 30.48 × 45.72 cm, Exoterra, Mansfield, MA) were lined with sphagnum moss, leaf litter, live philodendrons, a climbing log, and film canisters for egg laying and tadpole deposition. Frogs were kept on a 12:12 h light cycle and fed wingless *Drosophila melanogaster* fruit flies dusted with vitamin supplements three times weekly.

Pair bonds were established by cohabitating male-female dyads and allowing them to breed. Pair bond establishment was marked by the first successful reproductive bout consisting of tadpole deposition into a breeding pool. Once bonded, partnership longevity was calculated as the number of days from pair bond establishment to testing (132-1238 days across pairs). Tadpole deposition of each pair was monitored and recorded 3 times weekly. The lifetime reproductive output of pairs, calculated as the cumulative number of tadpoles deposited from partner establishment to testing (3-87 across pairs), was determined.

### Experimental design, and behavioral and hormonal sampling

We adapted an “empathy assay” for amphibians based on published assays used for rodents (13). We focused on males because time and availability of frogs during the COVID-19 pandemic did not permit studying both sexes, and we sought to complement other ongoing studies. Prior to trials, pair bonded males and either their female partner or a female non-partner were co-housed in a behavioral observation arena for 3 days to familiarize non-partnered dyads and acclimate. Behavioral observation terraria (36.4 X 21 X 12.4 cm) were lined with moist sphagnum moss, contained two canisters for shelter, and were covered with a transparent lid that was punctured to allow for ventilation and video recording using an aerially mounted GoPro camera, version Hero 6.

Following acclimation, trials were conducted from ∼13-17:30 hr. Male-female dyads were video recorded for pre-treatment behavior for 15 minutes. Following, dyads were sampled for pre-treatment corticosterone levels. In amphibians, corticosterone is released under basal/baseline conditions and peaks in response to stress, making it a physiological biomarker of the animal’s state along the relaxed-stressed dimensional axis (17–20). Amphibian corticosterone levels also change in response to social stimuli (21). These attributes make corticosterone a promising biomarker for the physiology of stress contagion between partners. To measure corticosterone, we used a pre-established water-born collection method, which is non-invasive and yields a valid representation of plasma values, including in amphibians (18,22–25). Briefly, each animal was placed in an isolated sterile petri dish (8 cm diameter X 2 cm high) containing 20 mL milli-q water conditioned with essential ions for amphibians (Josh’s Frogs RO R/x, Osowo, MI, USA) for 1 hour. To ensure that frogs remained submerged in the water, we covered the dish with a lid and used enough water volume to prevent frogs from residing on the sides of the dish above the water’s surface. Both frogs were then placed back into the behavioral terraria for 1 hour to recover from potential handling stress. Following recovery, females were removed and underwent 15 minutes of either a leg restraint stress treatment or a non-stress treatment. The leg restraint treatment, which involves gently restraining a hind leg to inhibit movement, is standard for inducing a mild acute stress for studying stress hormones in amphibians (26–29). In the non-stress treatment, females were placed into terraria like the behavioral observational terraria except enriched with leaves, a shelter log, and live philodendrons. In studies on stress contagion, it is important to rule out the possibility that state matching arises from observers responding to the stressor that demonstrators experience as a perceived threat to themselves, rather than responding to the demonstrator’s stressed emotional state (30). The advised way to do so in non-human animals is that the observer cannot perceive the stimulus that triggers the demonstrator’s emotional response (13,30). To control for this confounding source of firsthand stress in observer males, female treatments were unobserved by males. Treated females were then immediately reunited with males and behavior and hormones were re-sampled as described above (**Fig. 1**).

**Figure 1.**
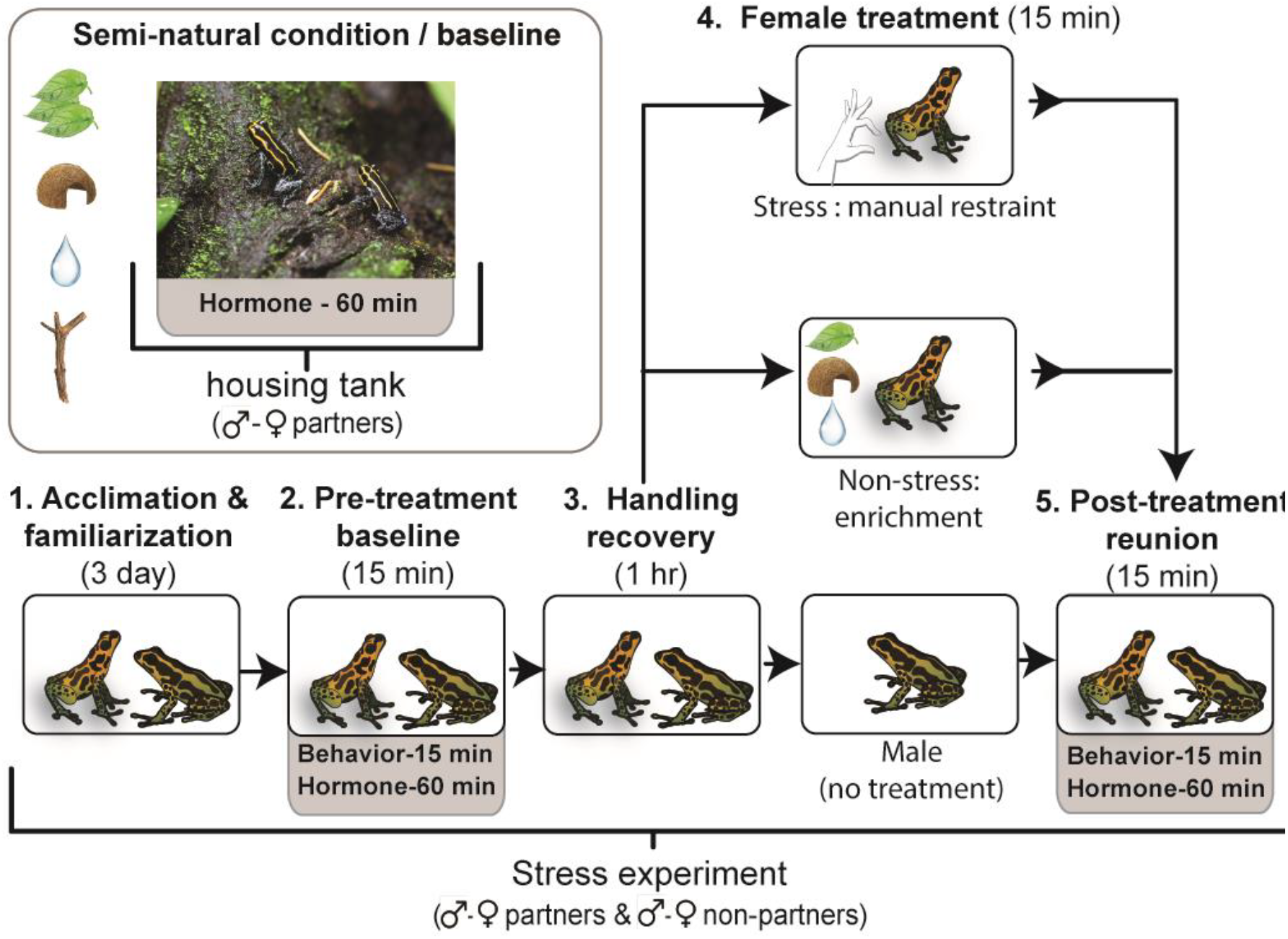
Design for testing partner-selective emotional contagion in pair bonded male *R. imitator* poison frogs. Pair bonded males were assayed for state matching the behavior and corticosterone level of partner females that underwent a stress treatment that males did not observe or experience themselves. To examine partner-specificity, each male was assayed with their female partner and a familiar female non-partner of similar reproductive salience. To better interpret these experimental results and establish the semi-natural (baseline) corticosterone state of pairs, we also examined corticosterone levels and matching of experimentally naïve pairs within their housing terraria. Picture is of a pair bond within its housing terrarium, taken by Daniel Shaykevich.

Experimental males (n = 9) underwent the aforementioned trials with each stimulus female (partner and non-partner), both before (pre) and after (post) each treatment (stress and non-stress). Each male’s female partner and non-partner were matched for social and reproductive status (all pair bonded and reproductively mature) and size (length: ± 1 g, weight: ± 0.6 mm) as best possible. The order in which males underwent female partner/non-partner and female stress/non-stress treatments was assigned arbitrarily and was separated by at least 1 day. Yet, we included the order of the trial as a random factor in subsequent analyses (see below). Immediately after the aforementioned trials, animals were placed back into their housing terraria. To control for circadian patterning of corticosterone associated with foraging and reproduction (17,18), we assayed for corticosterone within the same restricted time periods (approx.13:15-14:30 and approx.16:00-17:15 hr), and animals were food starved at least 17 hrs prior to and throughout testing. To compare individual corticosterone and male-female corticosterone co-variation between experimental and “semi-natural” baseline conditions, we also sampled corticosterone from pair bonded dyads that were experimentally naïve from within their housing terraria (n = 10). For housed animals, we sought to minimize the effects of circadian variation in corticosterone associated with feeding and reproductive activity by assaying for corticosterone between 14:00-17:00 hr and feeding the same quantity of food within the same 2 hr period. To control for the potential influence of life stage on corticosterone levels (31), all animals were mature adults. Corticosterone samples that did not meet quality control standards (see *Corticosterone quantification* section below) were removed from the study, resulting n = 10 housed dyads and the following dyads per experimental group: pre-experiment partner: n = 17, pre-experiment non-partner: n = 13, post stress partner: n = 6, post stress non-partner: n = 9, post non-stress partner: n = 5, post non-stress non-partner: n = 6.

### Behavioral quantification

All data were scored from video recordings by the same trained researcher, who was unaware of the experimental group. We used BORIS v7.12.2 software (32) to manually score the following behaviors: the duration and bouts of gaze towards conspecific, the duration and bouts of conspecific approach, the duration and bouts of freezing, the duration of refuge use, and the bouts of activity, joint refuge use, and coordinated freezing (see Supplementary Table 1 for behavioral ethogram).

To complement manual scoring, we also quantified the following behaviors using Annolid V.1.1.2 automated behavioral tracking and analysis software (33,34): average speed, total distance traveled, percentage of space used, and distance from conspecific (see Supplementary Table 1 for behavioral ethogram). To score behaviors in Annolid, we trained models to identify and track each individual within each of the 38 generated 15-minute videos, resulting in 37 unique models (one model per video, except for one model that successfully tracked 2 videos of the same male-female dyad). Specifically, each individual within the video was manually labeled across 20-134 key video frames. Models were then trained on the labelled frames with the initial weights based on the R50-FPN COCO segmentation Mask R-CNN model in Detectron2 (35) model zoo. The model’s accuracy in identifying individuals was then tested on the corresponding video and considered acceptable if it contained no more than 5 instances of misidentification that resulted in no more than 30 seconds of misidentification cumulatively. If the model failed this quality control standard, then it was re-trained by adding additional labeled frames.

### Behavior statistical analysis

All statistical analyses and data visualizations were performed in R (V. 4.0.3; the R Foundation for Statistical Computing). First, to account for and eliminate intra-individual sources of variation, we subtracted the pre-treatment baseline from the post-stress/non-stress treatment conditions. Then, we extracted behaviors, namely activity level, freezing duration, partner approach duration, partner gaze duration, total distance moved, and total area moved. Next, we minimized redundancy of the extracted behaviors using a varimax normalized principal components analysis (PCA) for each dyad type, using the function ‘principal’ within the *psych* package (36). We tested whether subjecting females to the stress treatment affected the individual or pairwise behavior of female demonstrators and/or male observers, using simple linear models. For this, we performed each model with each of the rendered components as response variables, and experimental conditions (post-stress/-non-stress), sex, and trial order as predictors. We further computed estimated marginal means (least-squares means) contrasts among conditions and sex, using Tukey’s adjustment method. To determine whether males displayed behavioral indicators of partner-selective emotional contagion, we examined male-female state-matching for stress related individual behaviors within the rendered principal components. For this, we ran simple correlations for each of the four experimental condition*dyad type combinations.

To test pairwise behaviors of dyads, we began by subtracting pre-experimental baseline values from post stress/non-stress values, to eliminate intra-individual sources of variation. Due to resulting zero-inflated data, we then conducted a two-part model following (37). Briefly, we created two new variables from each original zero-inflated pairwise behavior: 1) a binary variable that indicated whether an observation was zero or non-zero, which was used as a response variable in logistic regression models; and 2) a continuous variable that contained only the non-zero values (zeroes were replaced with NAs), which was used as the response variable in permutation tests. In models, we used the interaction between experimental condition (post-stress/-non-stress), dyad type (partner/non-partner), and trial order as predictors. Because the male-female proximity data contained a large amount of truly missing values, these data were omitted from the former analysis and analyzed separately for pre-treatment baseline and post-treatment condition. In each analysis, we used again a permutation test, with male-female proximity as response variable, and the interaction between experimental condition (stress/non-stress), dyad type, and trial order as predictors.

### Corticosterone quantification

Water-borne hormone sampling and extraction was conducted following the methods of (23–25). Briefly, water from the water bath was immediately stored at -80°C for up to 2 weeks. Following, samples were thawed on ice, and water was collected with a 20 mL sterile syringe and pumped through a C18 cartridge (SPE, Sep-Pak C18 Plus, 360 mg Sorbent, 55–105 μm particle size, #WAT020515, Waters corp., Milford, MA) at a rate of ca. 10 mL/min. Cartridges were then eluted with 4 mL of 96% EtOH into 8 mL borosilicate vials. Two mL was used for corticosterone analysis, and the remaining 2 mL was stored at -80°C for re-running samples that did not initially pass quality control or for future studies. Eluted corticosterone samples were then dried down with nitrogen gas at 37°C, resuspended with 250 µL of assay buffer (provided in ELISA kit, see below), and incubated overnight at 4°C. Reconstituted samples were brought to room temperature and shaken at 500 rpm for 1 hr.

A commercial enzymatic immunoassay was used to estimate corticosterone concentration (Enzo corticosterone ELISA kit, Catalog # ADI-901-097). Assay cross-reactivity with a related hormone, deoxycorticosterone, is 28.5%, and with other steroid compounds is < 2%, resulting in high specificity to corticosterone (see product manual for details). Samples were plated in technical duplicate and assays were performed following the manufacturer’s protocol. Plates were read at 405 nm, with correction between 570 and 590 nm, using a microplate reader (Synergy H1, BioTek Instruments, Winooski, VT, USA), and the concentration of corticosterone was calculated using a four-parameter logistic curve in the software Gen5 (version 3.05, BioTek Instruments, Winooski, VT, USA). The detection limit for the assay is 27 pg/mL, and samples that fell out of this range were removed from analysis. Samples with the average intra-assay coefficient of variation (CV) above 20% were also excluded from analysis.

### Corticosterone statistical analysis

All statistical analyses and data visualizations were performed in R (V. 4.0.3; the R Foundation for Statistical Computing). Prior to analysis, corticosterone concentration was log-transformed to achieve normal distribution when necessary. For all models, we conducted standard model diagnostics and assessed for outliers and deviations across quantiles using the ‘DHARMa’ package (38). First, we tested corticosterone state matching of male-female dyads within and across conditions. We ran analyses separately for each condition (housing tank, experimental pre-treatment baseline, and post-stress/non-stress treatment) and dyad type (partner and non-partner). Since corticosterone concentration for the housing condition was normally distributed, we performed a simple linear correlation using Pearson’s correlation coefficient. For the experimental conditions, we performed a linear mixed model LMM with male corticosterone log-concentration as the response variable, female corticosterone log-concentration and post-treatment condition (post-stress or post-non-stress) as fixed predictors, and frog identity and trial order as random factors.

Next, we tested whether male-female state matching predicted the longevity or lifetime reproductive output of partnerships. We ran a separate multiple regression for each condition (housing, experimental pre-treatment, post-stress, and post-non-stress treatments), using male corticosterone levels as the response, and female corticosterone levels, partnership endurance, reproductive output and trial order as predictors.

We tested for differences in corticosterone concentration between housing and experimental pre-treatment (baseline) condition. For this, we performed a linear mixed-effects model (LMM) using the ‘lmer’ function within the *lme4* package (39). We included log-transformed corticosterone concentration as the response variable, and the interaction between condition (housing and experimental pre-treatment baseline), dyad type (partner and non-partner), sex, and trial order as predictors, and the frog identification as a random factor. We further computed estimated marginal means (least-squares means) contrasts using the ‘emmeans’ function within the *emmeans* package (40). P-values were adjusted using Tukey’s method. Only pair bonded dyads were examined in the housing condition since animals were housed only with partners.

Finally, differences in corticosterone concentration between experimental treatment conditions were examined separately for each dyad type and sex also with LMMs. The models included log-transformed corticosterone concentration as the response variable, the interaction between treatment condition (non-stress and stress) and trial (pre- and post-treatment) as fixed effects, and frog identity and trial order as random factors. Post-hoc analyses were conducted using estimated marginal means with Tukey’s adjustment method.

## Results

### Males selectively state match their partners for corticosterone irrespective of the endurance or lifetime reproductive output of partnerships

Male corticosterone levels positively correlated with those of female partners within the housing baseline (Pearson’s correlation test: t = 2.69, *r*_*(8)*_ = 0.69, *P* = 0.02; **Fig. 2A**) and experimental pre-treatment baseline conditions (LMM: *β* = 0.5, t = 5.2, *R*^*2*^ = 0.642, *P* < 0.001; **Fig. 2B**). There was a similar trend after controlling for the post-treatment conditions (stress/non-stress), although to a statistically non-significant extent (*β* = 0.001, t = 2.33, *R*^*2*^ = 0.83, *P* = 0.06; **Fig. 2C**), likely due to small sample size (n = 5 and 6, respectively) and increased response variation (see Supplementary Tables 2 and 3 for details). Similarly, complementary research in voles has also discovered large effect sized, despite limited sample size (13). By contrast, male corticosterone levels did not correlate with those of female non-partners across any condition (pre-treatment: LMM: *β* = 0.2, t = 0.72, *R*^*2*^ = 0.041, *P* = 0.48; post stress/non-stress: LMM: *β* = -0.0007, t = -0.28, *R*^*2*^ = 0.74, *P* = 0.78; **Fig. 2D, E;** see Supplementary Tables 2 and 3 for details). The extent of male-to-female partner corticosterone matching was not explained by the endurance or lifetime reproductive output of partnerships (*P*-values for all treatment conditions ≥ 0.05; **Fig. 3**; see Supplementary Table 4 for details).

**Figure 2.**
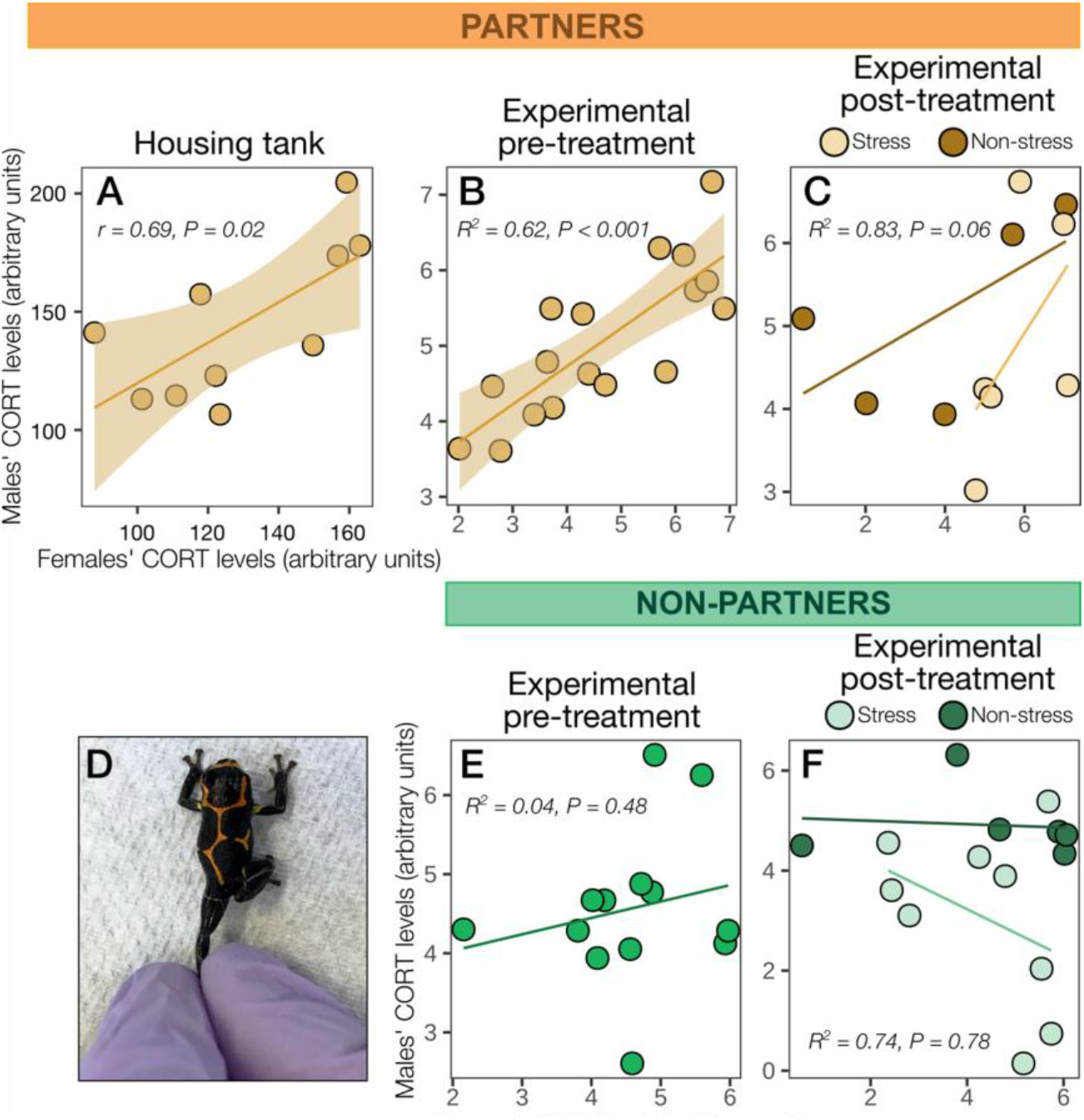
Partner-selective corticosterone matching in pair bonded *R. imitator*. Male corticosterone (CORT) level matched (**A-C**) female partners, but not (**E-F**) familiar female non-partners, across semi-natural and experimental conditions. Regression lines are shown, with shaded regions denoting 95% confidence intervals for statistically significant results. (**D**) Example of gentle leg-restraint stress treatment.

**Figure 3.**
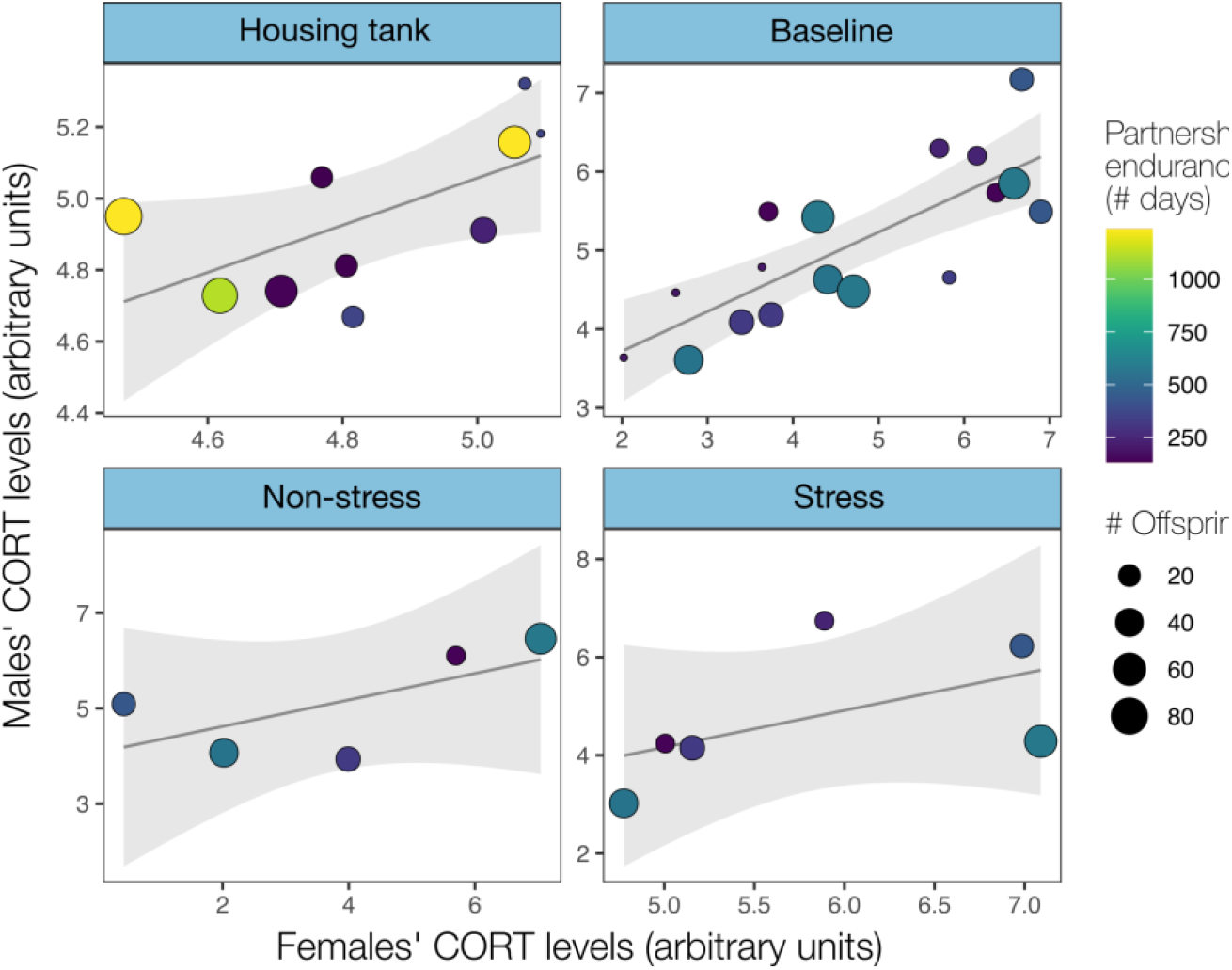
Male-female corticosterone state matching is irrespective of the endurance or lifetime reproductive output of partnerships. Shown are multiple regression plots of male-female corticosterone relationship across conditions after controlling for the endurance and total number of offspring of partnerships. Shaded area around multiple regression lines denote 95% confidence intervals.

### Stress contagion assay does not provoke a stress response, nor do males behaviorally state match females

Subjecting females to the stress treatment caused no increase in corticosterone levels in female demonstrators or male observers across time points, irrespective of whether they were normalized to baseline (planned least-squares means contrast: *P* ≥ 0.05 in all groups; Supplementary Tables 5, 6 and Supplementary Figure 1A, B, D for details). Principal component analyses for each dyad type (partners and non-partners) yielded three components that explained more than 70% of the total variance (See Supplementary Table 7). Subjecting females to the stress treatment had no effect on the individual or coordinated stress behavior of female demonstrators or male observers of partner and non-partner dyads in any time point comparison, irrespective of whether responses were normalized to baseline (*P* ≥ 0.05 for all treatment comparisons; Supplementary Tables 8-11 and Supplementary Figures 1C, E and 2D-G for details). Together, this indicates that the treatment did not provoke a stress response as intended. Finally, males did not behaviorally state match females in any experimental condition, except for gaze and approach towards non-partner females in the non-stress condition (*R*^*2*^ = 0.83, S = 20.32; *P* = 0.006), and freezing towards non-partner females in the stress condition (*R*^*2*^ = 0.67, S = 2.41; *P* = 0.04; Supplementary Table 12 and Supplementary Figure 2A-C for details).

## Discussion

### Hormonal state matching in poison frogs: partial evidence for emotional contagion

We discovered that male pair bonded poison frogs display hormonal state matching exclusively with female partners compared to familiar non-partner females. Specifically, males exhibited variation in corticosterone state, which was positively and exclusively correlated with female partners, both while cohabitating and experimentally, similar to findings in humans (41–45). Here, we discuss this as evidence for emotional contagion in an amphibian and suggest future clarifying experiments.

Emotional contagion is the process through which an individual’s emotional state (e.g., fear, joy, relaxed) comes to resemble or “match” that of another individual through observing their state (30,46,47). Since emotions are internal experiences that cannot be directly expressed beyond self-reporting, measuring emotional contagion in non-human animals relies on state matching of emotional indicators—the associated neural and physiological responses, and the downstream behavioral outputs that they motivate (30,46). Historically, behavioral indicators have been primarily used to infer emotional contagion. However, they are not a necessary indicator, since emotions are internal experiences that don’t always produce downstream behavioral outputs (2,30,46–48). Moreover, it is argued that none of these three types of indicators (neural, physiological, or behavioral) provide sufficient empirical evidence in isolation, because each is subject to pleiotropy (e.g., a given brain region is rarely activated by only one emotional state, a given hormone rarely serves only one function, or a matched behavior may derive from a natural tendency to imitate the movement of others in the absence of shared emotion) (30,46,47).

Therefore, the most robust reverse inference of emotional contagion is thought to derive from state-matching across multiple indicators, generating an overall pattern of readouts that helps contextualize interpretation by being more likely to be uniquely attributed to a particular emotional state (30,46,47). Moreover, it must be demonstrated that such state-matching is consequent of the observer directly responding to the demonstrator’s emotional state, as opposed to both subjects responding to an external stimulus in a similar manner (i.e., emotional convergence) (30,47). Among the few such studies that have been conducted in non-human animals that we are aware of (rodents (13), birds (14), and fishes (49,50)), all have found multi-modal state matching, strongly suggesting the presence of emotional contagion. For example, monogamous prairie voles match both the anxiety-related behavior and corticosterone increase of partners who have experienced an unobserved stressor (13).

We adopted the multi-modal approach, examining whether male observers physiologically and behaviorally state match female demonstrators that have been subjected to a stressor that they did not observe themselves. Experimentally, we discovered partner-selective physiological state-matching in males, such that male corticosterone state positively correlated with female partners but not familiar female non-partners. Inter-partner corticosterone state matching was also found in experimentally naïve pairs while cohabitating in their semi-natural housing terraria, suggesting that it is ecologically relevant rather than an experimental artifact. Similarly, sustained inter-partner cortisol co-variation has been repeatedly discovered in humans, where it is more pronounced between partners than non-partners (41) and occurs in both naturalistic and experimental conditions (42–45).

However, we found no evidence of coinciding behavioral state-matching. Although we examined a repertoire of behaviors relevant to stress and poison frog ecology (e.g., activity, freezing, refuge use, approach), possibly behavioral state-matching occurred in unexamined behaviors (e.g., toe tapping or jumping) (51,52). Alternatively, physiological state-matching might have occurred without behavioral state-matching. Without corroborating behavioral indicators, it is more challenging to interpret whether the inter-partner corticosterone state-matching observed represents an affective match consistent with emotional contagion, or alternatively the matching of other physiological processes that corticosterone pleiotropically serves in amphibians. Indeed, in addition to serving as the primary stress biomarker (53), corticosterone also mediates metamorphosis (31), and it follows a circadian rhythm in release that is associated with daily patterns of foraging and reproduction (17,18), which should conceivably be more similar between pair bonded individuals that share a foraging habitat and reproductive activity, respectively. The possibility of these developmental and diurnal autocorrelational effects can be ruled out since all animals were at the same developmental stage (reproductively mature adults) and assayed within the same restricted times of the day. The possibility of a shared foraging habitat effect can also be ruled out because feeding was standardized across housed partners and was abstained in experimental dyads prior to and during trials. A shared reproductive activity effect is also unlikely, since experimental trials occurred outside of peak reproductive hours in an observation arena void of reproductive environmental enrichment, and thus no reproductive activity was observed. Furthermore, if there was a shared reproductive activity effect, then we would expect male-female corticosterone matching to have occurred in both stimuli female treatments rather than to be exclusive to female partners, since both stimuli females had similar reproductive saliences—that is, both were familiar and acclimatized to males, reproductively experienced, and had similar morphometrics. Taken together, it appears unlikely that the observed partner-specific corticosterone state matching can be explained by a matching or synchronization of these alternative corticosterone-related processes.

The second finding that challenges the interpretation of emotional contagion is that inter-partner physiological state matching occurred in a static emotional state. Specifically, the stress treatment did not induce a detectable change in corticosterone or behavior in female demonstrators relative to either the pre-experimental baseline or semi-natural housing condition, to which males could co-respond in kind. Unlike other anurans (26–29), the leg restraint might be insufficient to stress *R. imitator* and thus the animals were consistently in a non-stressed or “relaxed” state. Alternatively, confinement in the water bath for 1 hr might have induced a maximum stress level in all hormonal treatment conditions, including baseline, resulting in animals consistently being in a stressed state. Regardless of the scenario, both relaxation and stress are valanced emotional states (54), which are subject to being contaged (47). Stable emotional states are also subject to being contaged (termed morphostatic covariation) (55). However, they can also be explained by emotional convergence (30,55). Such that, rather than physiological state-matching arising from males directly responding to the emotional state of females (i.e., the “transference” of emotion), it arose from both sexes directly responding to a shared environmental stimulus in a similar emotional manner. The possibility of emotional convergence can be ruled out by two observations. First, the stress treatment that females experienced was unobserved by males. Secondly, males experienced the same experimental paradigm and therefore environmental stimuli when assayed with each test female (partner and non-partner), yet their physiological state-matching is exclusive to female partners. The lack of matching with non-partner females further suggests that males are not simply responding to the emotional cues that female partners produce without regard for their identity (another form of emotional convergence), but rather to the emotional state of the female partners themselves (13,30). Finally, as inter-partner physiological state matching also occurs in the semi-natural housing environment, females may also display hormonal state matching, and this should be tested in the future.

In conclusion, it appears most likely that the partner-specific corticosterone matching displayed by males represents emotional contagion, such that it is a similar emotional state that is being transmitted directly and exclusively between partners. Socially selective emotional contagion requires emotional regulation, which was thought to rely on cortical brain regions and therefore be unique to mammals (56). However, evidence for it now also exists in amphibians (current study), fishes (50) and birds (14), suggesting it is phylogenetically widespread. This capacity to be affected by and share the emotional and physiological state of others has important implications for considering the wellbeing of non-human animals in captivity and nature (3). To test the emotional contagion hypothesis more conclusively in *R. imitator* however, further studies that elicit coinciding behavioral and neural indicators as well as a matched change in emotional state between partners (i.e., morphogenic covariation) (55) are now needed.

### Inter-partner physiological state matching is independent of the longevity or reproductive output of partnerships

In pair bonding species, emotional contagion is considered a mechanism that promotes cooperation, enhancing lifetime reproductive output and survival. In this process, partnership endurance might be a cause and/or consequence of emotional contagion. Indeed, several studies on pair bonding and bi-parental species have demonstrated partnership longevity increases bi-parental care and reproductive fitness (57,58). However, we found that inter-partner physiological state matching did not correlate with partnership longevity or lifetime reproductive output, indicating that it has no bearing on these attributes within this small sample size. Similarly, in human partnerships, cortisol co-variation can be only marginally linked to partnership duration (59). Alternatively, these relationships went undetected in the current study because they exist on a timescale that was not surveyed or were artifactually masked in captivity. Indeed, in wild *R. imitator*, pairs have not been observed to remain intact beyond one reproductive season (∼120 days) (16); whereas, in the captive colony used, partnerships were assayed after 132-1238 days of endurance, potentially leading to a “ceiling effect”. Additional work is needed to resolve these alternative possibilities to inform the potential biological significance of inter-partner physiological state matching in *R. imitator*.

## Conclusions

We sought to examine the empathetic-like phenotype for the first time in an amphibian. We found that pair bonded mimetic poison frog males display corticosterone, but not behavioral, state matching exclusively with female partners. This hormonal covariation cannot be explained by the confounds of autocorrelation, non-emotionally related functions, or emotional convergence. These results tentatively suggest emotional contagion in an amphibian, which along with similar findings in other taxa, indicates it is phylogenetically widespread. Paradigms that elicit coinciding neural and behavioral indicators and morphogenic co-variation are needed for further corroboration. Further studies on ancestral forms of empathy in non-mammalian vertebrates are warranted.

## Acknowledgements

We thank the O’Connell Lab for helpful feedback and Daniel Shaykevich for providing the picture of paired *R. imitator*. This project was funded by a New York Stem Cell grant to L.A.O. (grant NYSCF-R-NI58) and a Stanford Wu Tsai Institute Postdoctoral Fellowship to J.P.N. We acknowledge the animals that were used to conduct this study. We further acknowledge that this research was conducted at Stanford University, which resides on the ancestral and unceded land of the Muwekma Ohlone Tribe. L.A.O is a New York Stem Cell Foundation – Robertson Investigator.

## Author contributions

**Jessica Nowicki:** Conceptualization, Methodology, Investigation, Data curation, Writing – original draft, visualization, Project administration, Funding acquisition. **Camilo Rodríguez:** Methodology, Software, Validation, Formal analysis, Data curation, Writing - review and editing, Visualization. **Julia Lee:** Investigation, Writing – review and editing. **Billie Goolsby:** Validation, Investigation, Data curation, Writing – review and editing. **Chen Yang:** Methods, Software, Formal analysis, Resources, Data curation, Writing – review and editing. **Thomas Cleland:** Methods, Software, Writing – review and editing. **Lauren O’Connell:** Methodology, Resources, Writing – review and editing, Supervision, Project administration, Funding acquisition.

