## Supplemental material for "Physiological state matching in a pair bonded poison frog"

**Supplementary materials**

Jessica P. Nowicki^1^*, Camilo Rodríguez^1^, Julia C. Lee^1^, Billie C. Goolsby^1^, Chen Yang^2^, Thomas A. Cleland^2^, Lauren A. O’Connell^1^*

^1^Department of Biology, Stanford University, Stanford, California, USA

^2^Department of Psychology, Cornell University, Ithaca, New York, USA

*To whom correspondence should be addressed:

Department of Biology

Stanford University

371 Jane Stanford Way

Stanford, CA 94305

**Supplementary Table 1: Ethogram of *Ranitomeya imitator* behaviors quantified from video recordings taken during empathy assay.**

| **Behavior** | **Description** | **Scoring software** |
| --- | --- | --- |
| *Individual* |  |  |
| Gaze – bouts | Total number of times animal is facing towards other conspecific | BORIS |
| Gaze – duration | Total duration animal is facing towards other conspecific | BORIS |
| Approach – bouts | Total number of times animal moves towards other conspecific | BORIS |
| Approach – duration | Total duration of times animal moves towards other conspecific | BORIS |
| Freeze – bouts | Total number of times animal is motionless for at least 1 min. and completely visible | BORIS |
| Freeze – duration | Total duration of time animal is motionless for at least 1 min. and completely visible | BORIS |
| Refuge use – duration | Animal is attempting to be concealed within moss or in cup | BORIS |
| Activity – bouts | Total number of times animal is in motion, either stationary or through space | BORIS |
| Activity – speed | Average speed animal has moved through space | Annolid |
| Activity – distance | Total distance animal has moved | Annolid |
| Space use | Total area of space that the animal used (cm2) | Annolid |
| *Male-female dyad* |  |  |
| Male-female proximity | The average distance between animals | Annolid |
| Male-female joint refuge use | Both animals are refuging together in moss or cup | BORIS |
| Coordinated freezing | Both animals are freezing at same time | BORIS |

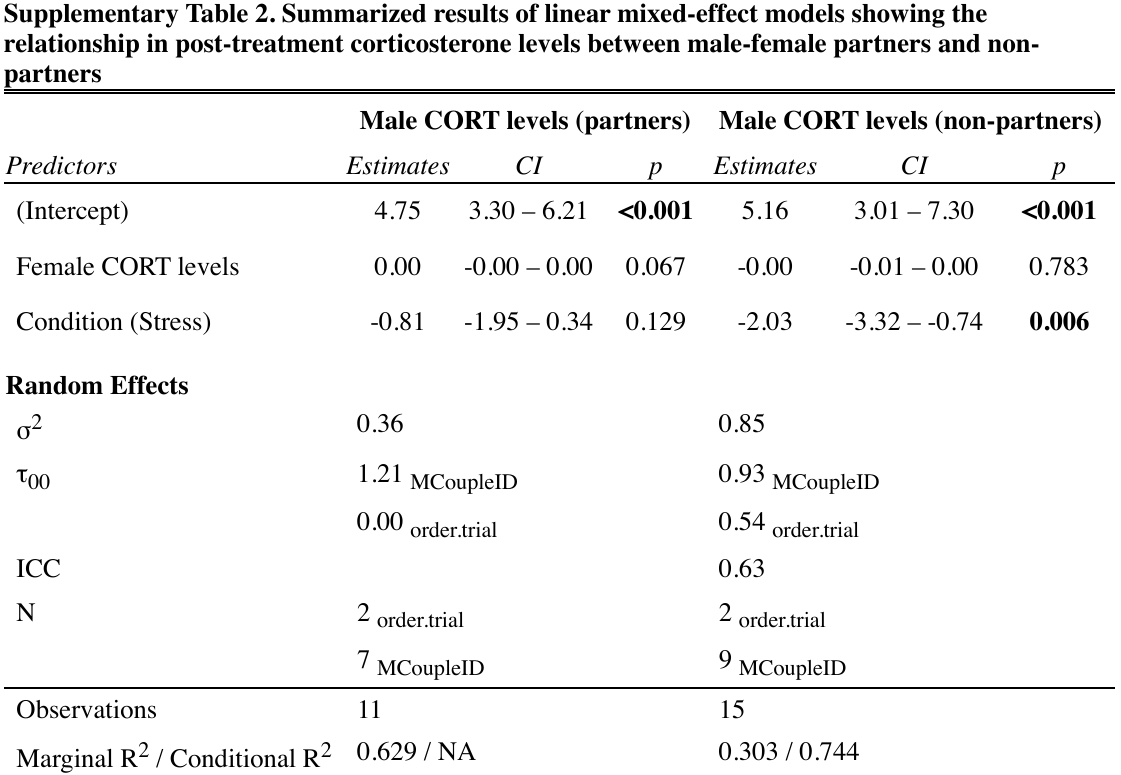

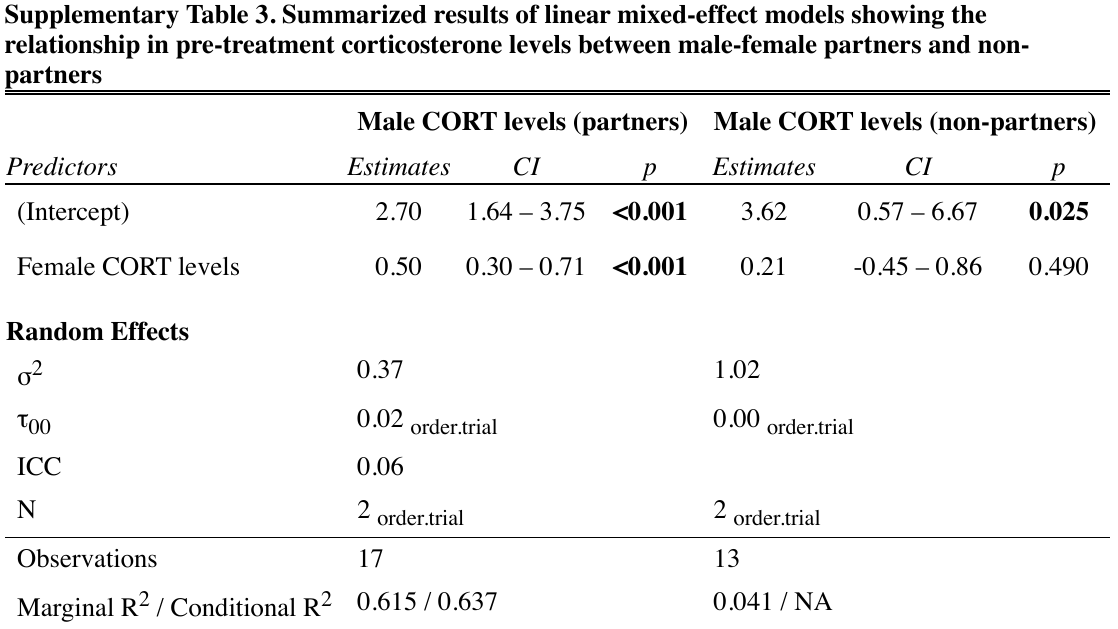

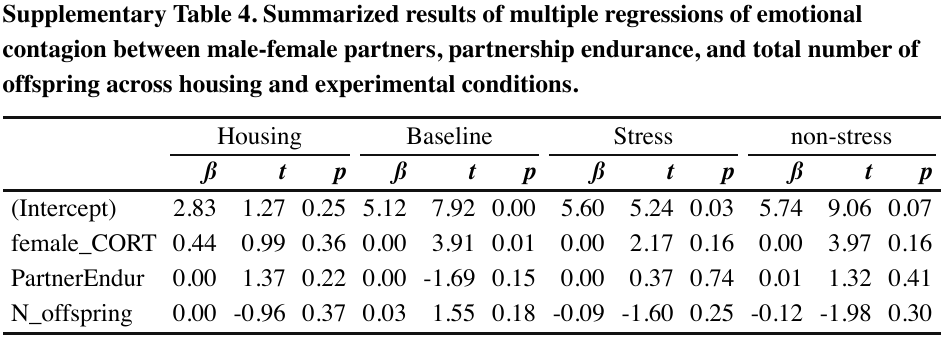

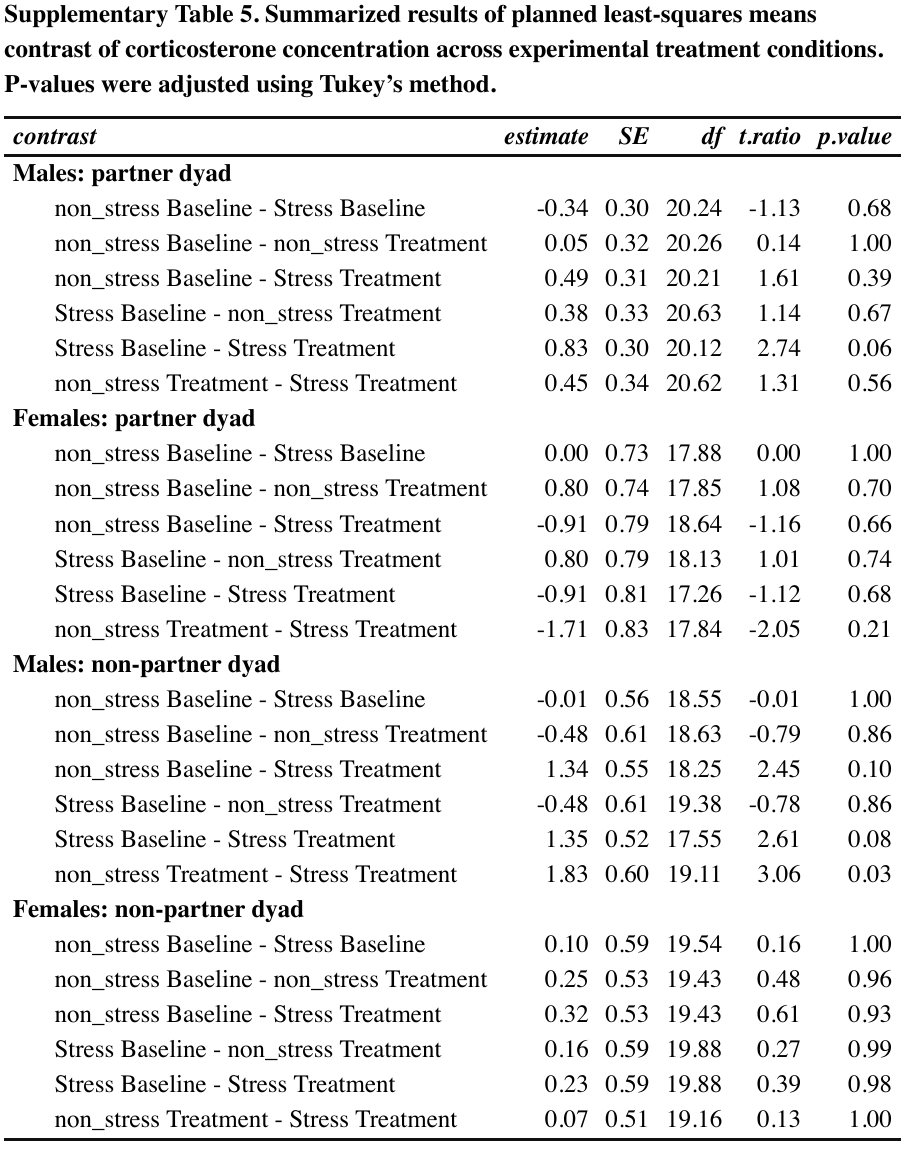

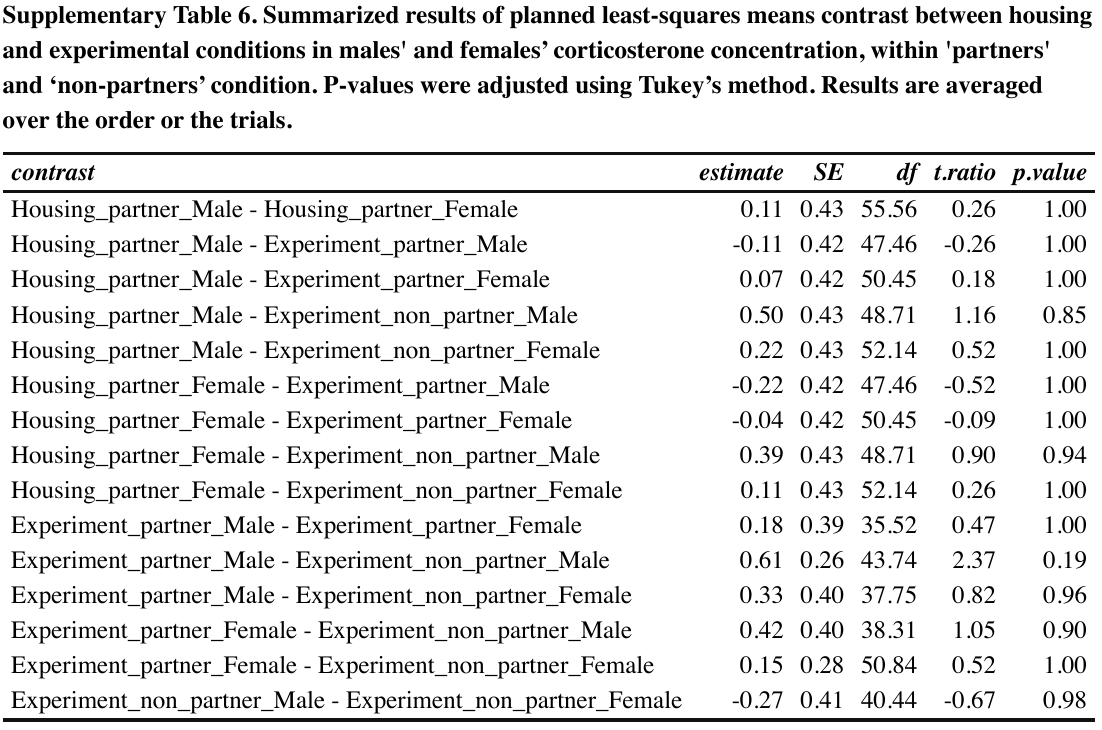

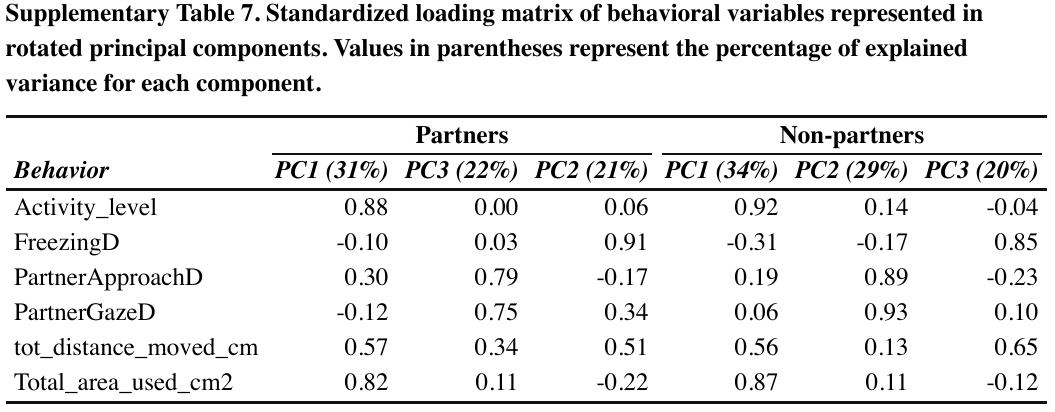

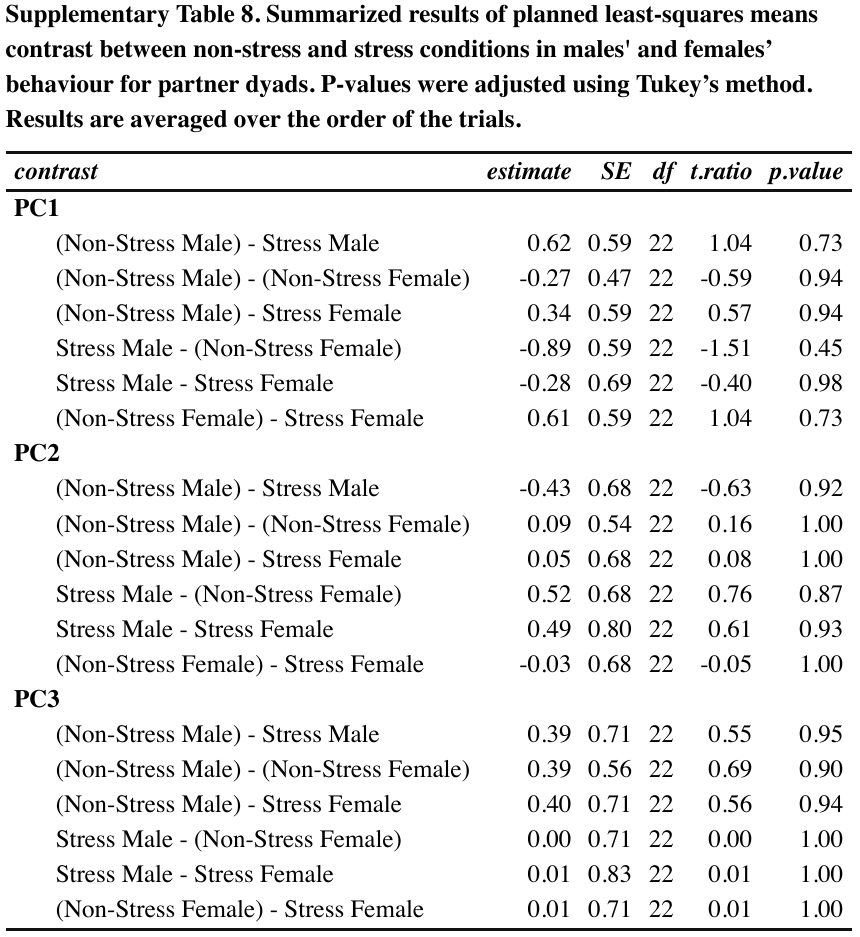

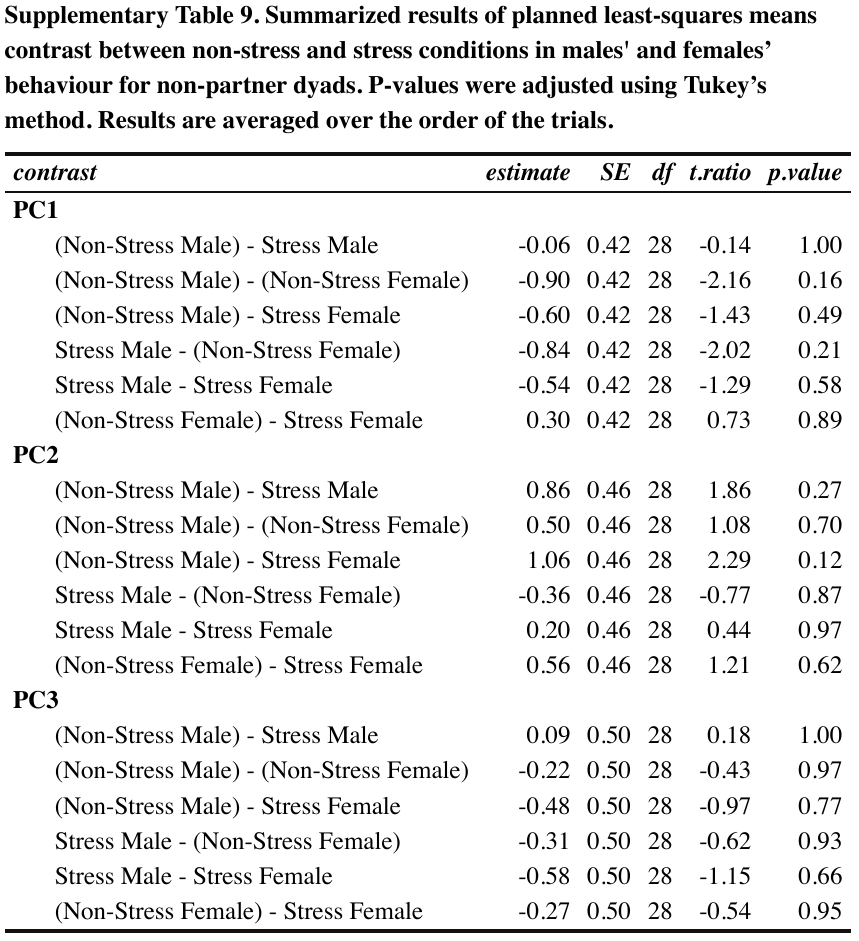

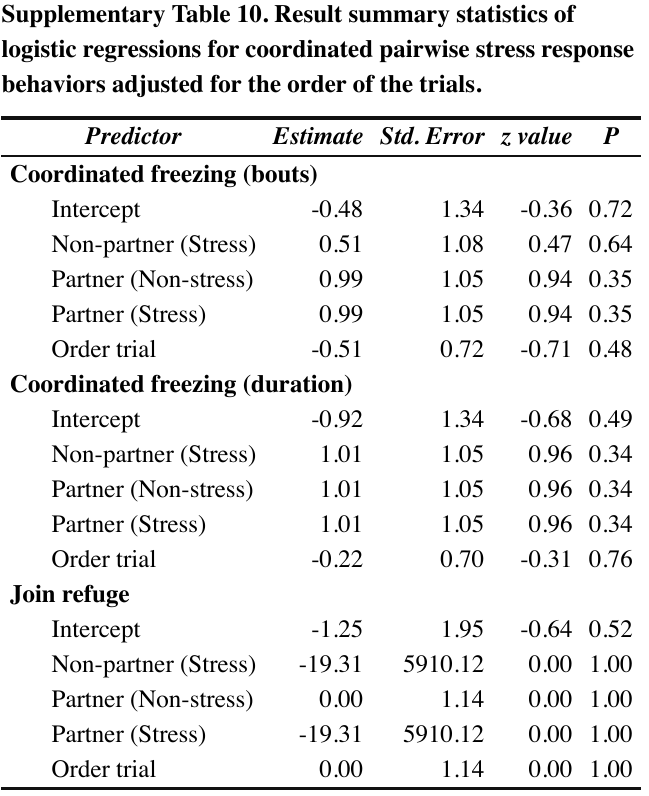

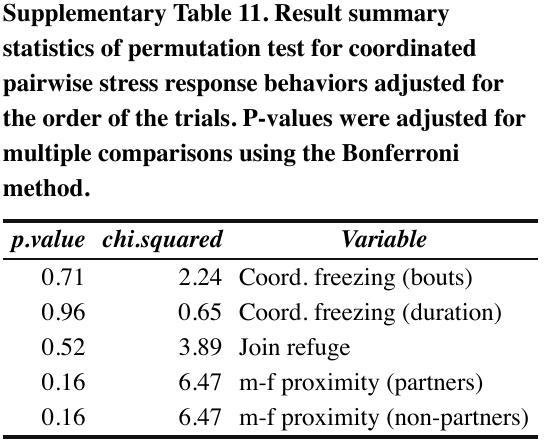

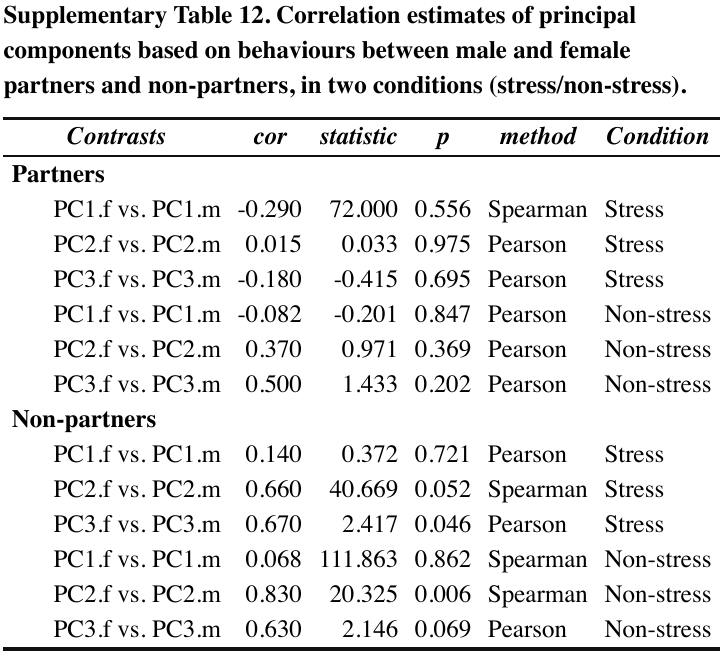

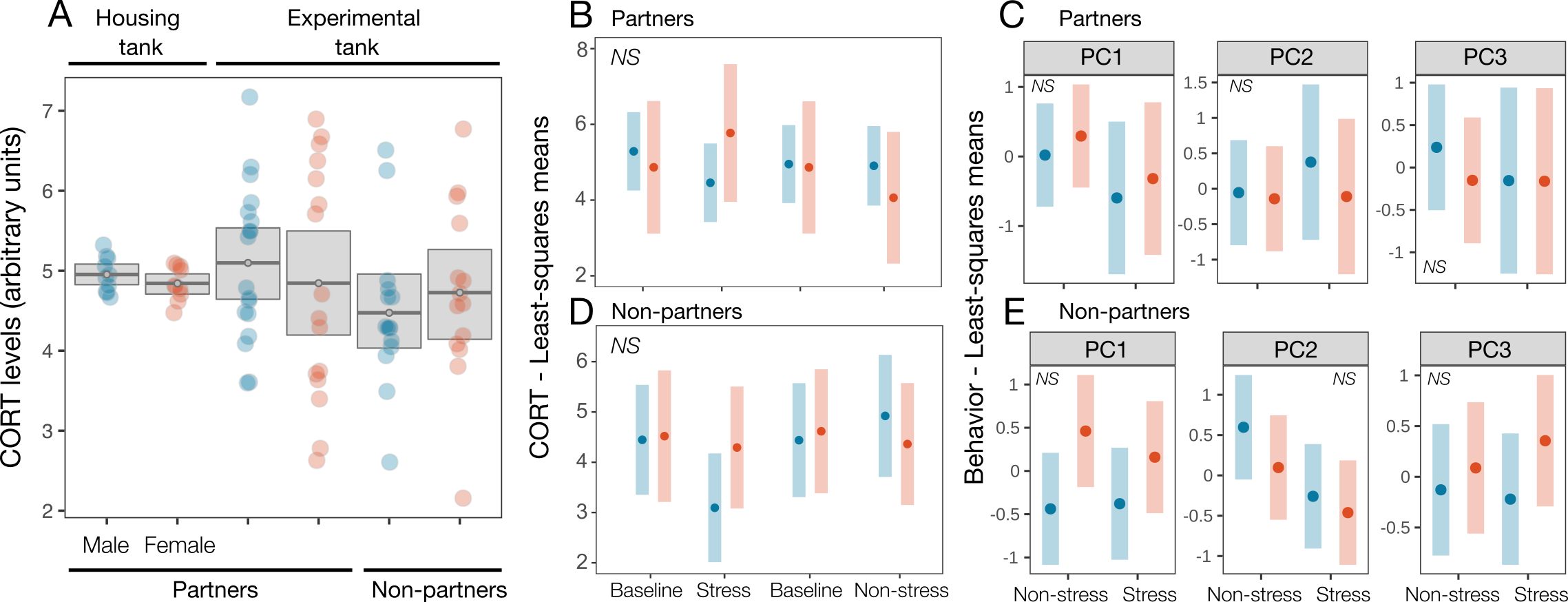

**Supplementary Figure 1. Subjecting females to a leg restraint treatment did not cause a corticosterone (CORT) or behavioral stress response in either female demonstrators or male observers in partnered or non-partner dyads.** **(A)** Water-borne CORT concentration does not differ between conditions (housing vs. experimental pre-treatment (baseline)), dyad (partner vs. non-partner), or sex. Dots and horizontal bars represent the mean, and grey boxes represent 95% confidence intervals. **(B, D)** Neither water-borne CORT concentration nor **(C, E)** behavior differs between experimental conditions (pre-treatment baseline vs. post non-stress/stress treatment), in either dyad or sex. Nine behavioral types are represented by 3 non-redundant principal component categories: PC1 = male-female interaction, PC2 = activity, PC3 = stillness. Behavioral values shown are relative to the experimental pre-treatment (baseline) condition. **Panels** **B-E**: dots represent the mean and bars represent 95% confidence intervals for the least-squares means. NS = non-significant (P ≥ 0.05).

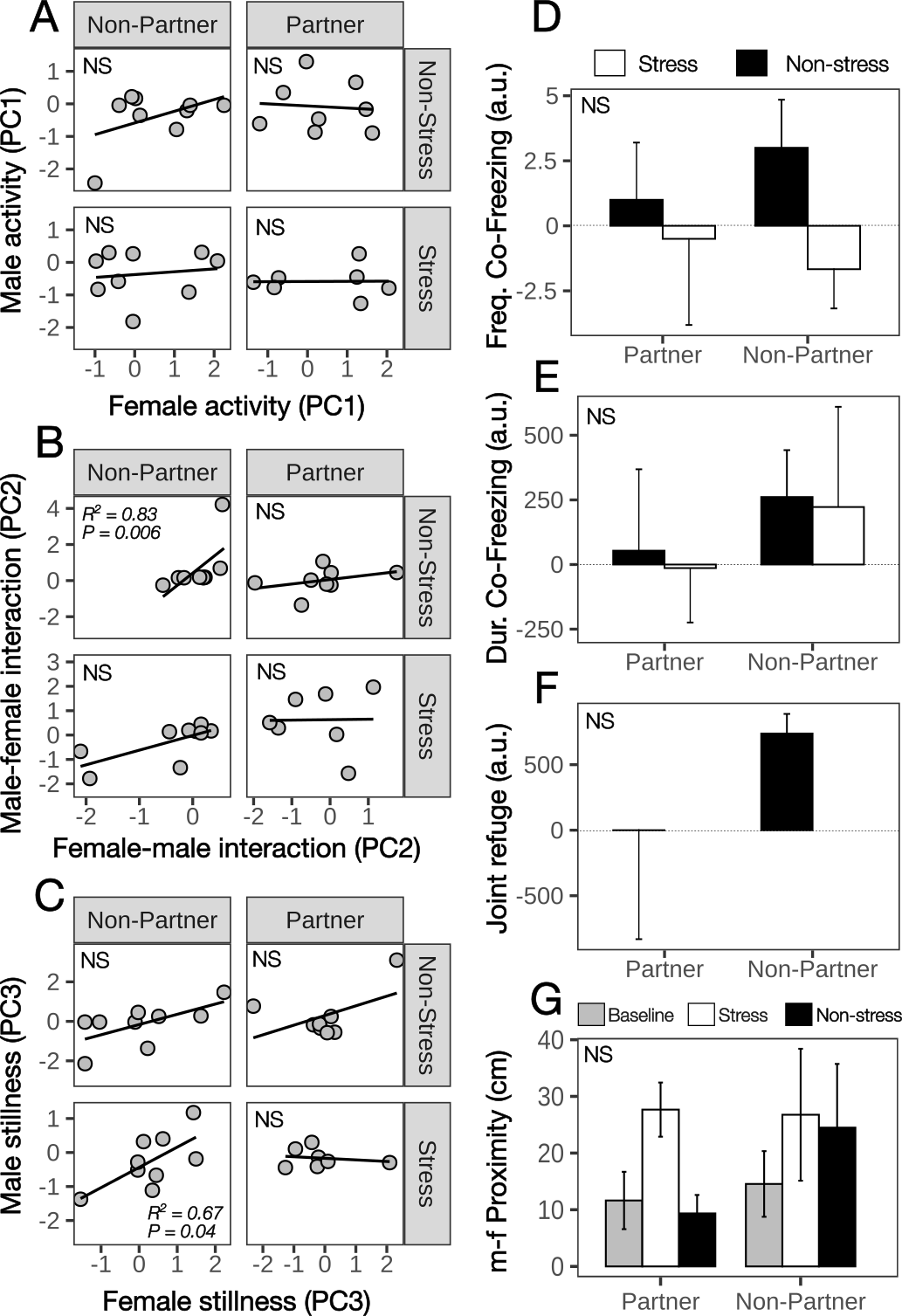

**Supplementary Figure 2. Overall, males do not behaviorally state match females, nor do they increase dyadic fear responses with females after females are subjected to a stressor treatment.** **(A-C)** Overall, males do not behaviorally state match female partners or non-partners in any condition. Shown are male-female correlations of 9 behaviors reduced to 3 non-redundant principal component categories: **(A)** male-female interaction, **(B)** activity, and **(C)** stillness levels, after subtracting pre-experimental baseline levels. **(D-G)** Likewise, the stress treatment does not affect coordinated dyadic fear response behaviors relative to experimental pre-treatment baseline or post- non-stress treatment conditions, in either partner or non-partner dyads. Values shown are relative to the experimental pre-treatment (baseline) condition, except for m-f proximity, which is an absolute value. Bars represent the mean and whiskers denote 95% confidence intervals. NS = non-significant (P ≥ 0.05).
